# Pretend Comprehension Enhances Social and Exploratory Behaviors in Human Toddlers and Adults

**DOI:** 10.64898/2026.03.24.713388

**Authors:** Camilo Gouet, Cristina Jara, Cristóbal Moenne, Damaris Collao, Marcela Peña

**Affiliations:** Laboratorio de Neurociencias Cognitivas, Escuela de Psicología, Pontificia Universidad Católica de Chile, Avenida Vicuña Mackenna 4860, Macul, Santiago, Chile; National Center for Artificial Intelligence CENIA, Chile, FB210017, Basal ANID; Departamento de Fonoaudiología, Facultad de Medicina, Universidad de Chile; Departamento de Kinesiología, Escuela de Ciencias de la Salud, Facultad de Medicina, Pontificia Universidad Católica de Chile

**Keywords:** Pretend play, symbolic play, Theory of Mind, social development, visual exploration, curiosity, information seeking

## Abstract

Pretend play is a hallmark behavior in childhood where children create nonliteral meanings. Empirical data supporting the role of social cognition and the decoupling from literality are still scarce during early development. Here we examined how the comprehension of pretense modulates visual exploration in 19-month-old toddlers (n = 44) and adults (n = 65) when they were exposed to short videos with an actress performing either real actions (e.g., eating jelly) or pretend actions (e.g., pretending to eat with imaginary food), while varying the complexity of those actions. We analyzed participants’ exploration of the actress’s face as exploitation of social information. We showed that all observers paid more attention to the face in pretend episodes than in real ones, measured as longer total looking time in adults and more fixations and revisits to the face in both age groups. We also found more gaze shifts (a measure of information sampling) between the face and the moving hand in pretense across age groups, mainly at the initial stages of the actions. Additionally, analyses of the scanpaths’ structure using gaze entropy revealed less order in the exploration of pretend movies at both ages. Both social and visual trajectory effects emerged mostly in complex pretense actions. Together, our results indicate that from its developmental origins, the comprehension of pretense relies on social inferential processes linked with information seeking and exploration.

## Introduction

Our human affinity for fiction begins early in life, in the form of a behavior called pretend play or symbolic play. During pretend games, objects and actions acquire nonliteral meanings^1^, as when children use a box to represent a battleship or play tea party with empty cups. Understanding episodes like these, while amusing, imposes a surprisingly non-trivial computational problem for observers to somehow override perceptual inputs filled with counterfactual or missing evidence and focus on meaning. How do children solve this? Do they read a partner’s intention to play behind these acts to guide their interpretation of actions, or do they grasp their meaning in purely behavioral codes? More generally, does the way observers gather and integrate visual information change in pretend contexts compared to real-world scenarios? The present study explores these questions in early development, when pretend play emerges.

Harris and colleagues pioneered experimental approaches to the study of pretense^2,3^, showing that the capacity to comprehend nonliteral episodes emerges at around 18 months. Subsequent studies employed looking time techniques, thus circumventing the need of a verbal or motor response on the part of subjects^4,5^, showing, for instance, that upon being familiarized to a pretense scenario with an actor pouring imaginary water into one of two glasses, 15-month-olds looked longer at the inconsistent (i.e. drinking from the ‘empty’ glass) compared to the consistent action (drinking from the ‘full’ glass), suggesting that they can grasp the goal of the action and expected the actor to act consistently with it. Notably, even though no actual liquid was involved. This capacity to comprehend and reason with pretend entities is a key developmental milestone in the second year of life, scaffolding more complex and extended forms of shared pretense and role play in toddlers and preschoolers^6^, which continue to develop into adolescence and adulthood^1^.

Theories of pretense comprehension posit that some form of decoupling must ensue that frames^7^ or ‘flags’^8^ a pretense situation as having a fictitious status distinct from reality. Studies with preschool children show indeed that they distinguish play from no-play situations, and differentiate classes of fictional worlds, e.g., so that what happens to Batman has no bearing on SpongeBob^9^. Then, to comprehend the actions composing the episodes, observers would need to recognize a partner’s intention to pretend and not to act literally, relying on inferential processes akin to belief and desire attribution, all belonging to theory of mind (TOM)^10^. This social mechanism, put forward by Alan Leslie and colleagues^10,11^, states that pretense and TOM abilities emerge together from the same conceptual system in the second year, an account that ignited a long-lasting and ongoing debate in the field^12–14^. Competing accounts have argued that mindreading capacities to metarepresent (i.e., to represent what others think or know as distinct from one-self’s representations) in pretense and TOM alike do not emerge in full-fledged form until around 4 years^15^, while further models have dismissed altogether such social underpinnings, maintaining that observers can understand make-believe actions as special ‘as-if’ behaviors^2,12^, without invoking social inferential mechanisms.

Empirical studies aimed at testing these hypotheses have given support to the social view of pretense in young children and adults, showing, for instance, that in 4- to 5-year-olds there are positive associations between theory of mind and pretend abilities^16,17^. Likewise, brain imaging studies in adults report that when exposed to short clips depicting either pretend or real actions (e.g., a person is shown pretending to drink versus drinking real juice), observers exhibit higher activity in medial prefrontal and temporo-parietal areas in pretense than in serious actions^18–20^, two regions involved in TOM tasks^21^, supporting an involvement of social inferences in adult observers. However, it is currently unclear if these findings hold in early pretense. One study^22^ reported no association in 18-month-olds between pretense comprehension and performance in a false-belief task (traditionally used to measure TOM), whereas another study^23^ found that performance in false-belief at 18-months predicts engagement in pretense at 4-5 years. Mainly because of its scarcity, these studies provide only fragmentary and indirect empirical evidence supporting theories at an early age.

In the current work, we aimed to contribute to filling this gap by evaluating the early steps of pretense comprehension in healthy toddlers and comparing them with adults. We created a series of short movies with an adult performing either real-world or pretend actions and monitored the visual behavior (eye movements) while participants watched the movies. Our first goal was then to test the hypothesis that understanding pretense would engage social processes right from the second year. To do so, we compared the extent to which participants focused their attention to the performer’s face in both types of movies. Previous studies indicate that the face is a source of social information for observers^24^, which helps infer goals and intentions^25^, and emotional states^26^, through a mechanism of social gazing that is active from the first months of life^24,27^. We employed dynamic Area-of-Interest analysis^28^, a method that allows pinpointing, moment-to-moment, where attention is and how it fluctuates between visual regions via gaze shifts (rapid movements of the eyes). For adult observers, we expected to find more gaze shifts between the face and other regions in pretense than in real actions, consistent with previous studies supporting social processes in adulthood underlying pretense^18–20^. For toddlers, we anticipated that a similar pattern as in adults, i.e., the amount of social gazing to the face, will favor the early emerging social account on pretense^10,11^, whereas a lack of similarity will be more consistent with the protracted view of social development^15^.

Beyond scrutinizing the social underpinnings of pretense, the use of dynamic analyses allowed us to evaluate the overall organization of visual exploration, gaining insights into the decoupling process and play mode in pretense. As mentioned above, entering a play mode is key to keeping reality-fantasy separated, but it is also thought to foster exploration and flexible reasoning. Studies in 4-to-6-year-olds report that performance in syllogistic deductions^29^ and causal reasoning^30^ improves by framing tasks as pretense games, and behavioral studies in preschoolers have recently revealed unusual and more exploratory behaviors (e.g., to walk from one point to another or recover a stick) when goals were presented as play versus instrumental actions^31^.

Thus, as a second goal, we asked, if in addition to changing the way observers gather and process social information, pretense modulates the overall visual exploration pattern. Indeed, vision is conceived as an active process that samples visual information from the environment and integrates it over time^32^. Interestingly, previous reports show that trait curiosity modulates the way adults explore visual scenes^33^ and that curiosity is an inherent ability of young children^34^. Here, we studied the order within gaze trajectories using gaze entropy, a measure previously used to measure the level of structure of visual scanning in healthy adults and amnesic patients^34^. We expected to observe less organized paths in pretense than in serious (real) scenarios in both age groups, consistent with an enhancement of an exploratory mode in pretense.

We included two types of actions in the stimuli set that differed in the number of steps involved, categorized as ‘simple’ and ‘complex’ ones. Similar distinctions have been made in previous studies^2,3^, reporting differences in comprehension performance between types of actions in toddlers. The inclusion of varied actions contributes to the ecological validity of the findings we obtained, and we expected the hypothesized effects on social and exploratory behavior referred above to occur across actions.

In sum, our study looked for new empirical data to evaluate the putative social nature of pretense and whether visual exploration is enhanced in pretense episodes compared to real-world scenarios, suggesting curiosity-like exploration patterns during early development, similar to adults.

## Results

Building upon previous adult studies^18–20^, we created a series of short movies with an actress performing real-world, daily actions (e.g., eating jelly, drinking juice) and their pretend, pantomimed counterparts (pretending to eat, pretending to drink with imaginary liquid). Orthogonal to the real-pretense factor, the four actions presented in each condition (real, pretend) differed in the number of steps involved and/or the objects manipulated within them, as shown in **Figure 1A**. As in previous studies^2,3^, we refer to one-step actions (’*eating a cookie*’ and ‘*drinking juice*’) as ‘simple’ and two-step ones (’*serving and eating jelly*’ and ‘*serving and drinking juice*’) as ‘complex’.

**Figure 1:**
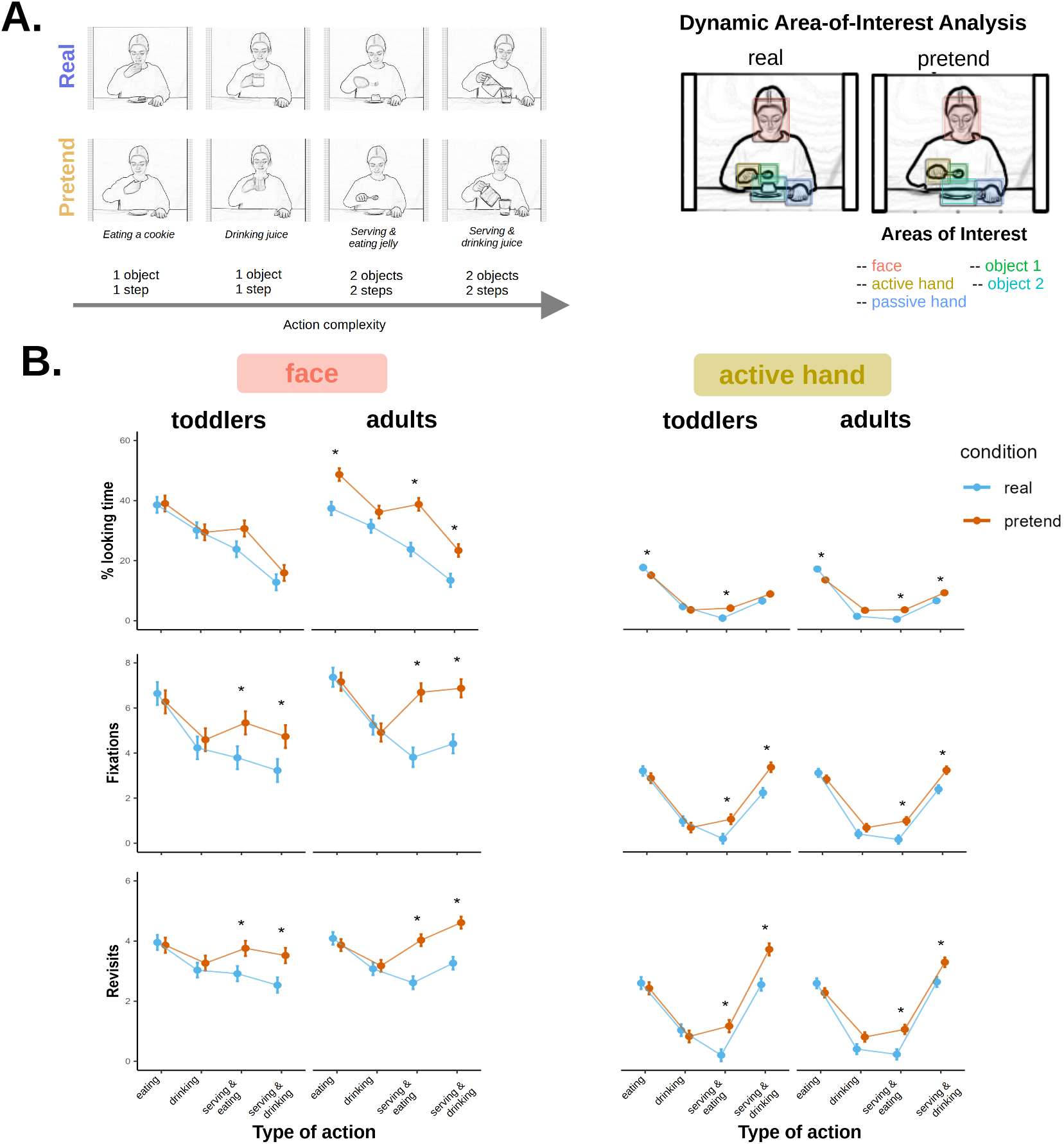
Video stimuli and dynamic area of interest analysis. **A**, snapshots of the four movies presented in each group, from simpler to more complex ones. During the actions, the actress approaches one hand to the central elements over the table to take some food or juice, then eats or drinks, and finally leaves the objects as located at the beginning (detailed in Figure 2A and Methods). The number of steps below each action’s name refers to those performed to take the food or juice. So, in ‘*eating a cookie*’ and ‘*drinking juice*’ the actress directly grabs a cookie or holds the cup with the active hand, in one step, whereas in ‘*serving and eating jelly*’ and ‘*serving and drinking juice*’ she uses a spoon to serve herself a piece of jelly or uses a jar to pour juice onto a glass. Right side, the areas-of-interests (AOIs) overlaid as colored boxes (red: face, yellow: active-hand, green: object 1 (located at the center: cup, plate or glass), cyan: object 2 (spoon or jar in complex actions), blue: passive-hand) for the action ‘*serving and eating jelly*’. **B**, outcomes of the dynamic AOI analysis across actions, for the face and active hand. The dots are estimated marginal means (EMM) across subjects for the measures dwell time, number of fixations, and number of revisits. Error vars correspond to the standard errors of the EMMs. *, p < 0.05.

We individually exposed 109 participants (toddlers n = 44, mean age = 19.56 months, age range = 16.26 - 20.00 months; females = 25; adults, n = 65, mean age = 20.55 years, age range = 19 - 24 years, females = 53) to both types of movies. Participants were allocated to start with either real or pretense conditions, and were instructed to passively watch the respective movies while their eye-movements were monitored using an eye-tracker (see details in Methods). The four actions per experimental condition (real & pretense), were presented two times each, alternately (average duration of movies = 8.11 s, range [6.17s −11.48s], see snapshots in Fig. 1A). As in previous studies in TOM with similar design^36^, here we focused the analyses in comparing the first experimental condition (real and pretend) that each group was first exposed to.

### Social nature of pretense comprehension

To test the hypothesis that social processes underlie pretense comprehension, we measured and compared observers’ attention to the face of the actress performing make-believe versus real actions. The dynamic Area-of-Interest (AOI) approach used (Fig. 1A, right side), helped us explore this hypothesis by analyzing the differences between participants’ visual behavior over areas with more or less role in social processing. We focused our analysis on looking time and dynamic measures targeting the face and the active hand (see example videos with the moving AOIs in SI). The former was the key target of our analyses, and the latter was evaluated as previous studies have shown that perception of the active hand contributes to action understanding in children and adults^28^.

First, we evaluated global patterns of attention through dwell time (% of total looking time to an AOI). As shown in **Figure 1B**, in both age groups, we observed a higher percentage of looking at the face in pretend than in real condition across actions. To statistically evaluate differences between experimental conditions (real versus pretend), we fitted a linear mixed-effects model of dwell time to the face as the dependent variable, with age and condition (between-subjects factors), type of action (within-subject factor) and their interactions as fixed factors and a random intercept for subjects. The goal of using this omnibus modeling approach was to obtain a measure of experimental error (non-explained variance) accounting for the mixed factorial design of the study using most available data. Thus, while we report the significance of the model’s parameters, the focus was on testing behavioral differences between real and pretend conditions to evaluate the social hypothesis.

Our results yield a main effect of condition (F(1,105) = 8.95, P = 0.003), type of action (F(3,713) = 222.70, P < 0.001), a marginal main effect for age (F(1,105) = 3.73, P = 0.056), and a significant two-way interaction between condition and type of action (F(3,713) = 7.26, P < 0.001). No further effects were significant (all Ps > 0.06), including the triple interaction between age, condition and type of action, which means that the way condition and type of actions interact is similar between age groups. This is visually apparent from the similar downward trends of dwell time with action complexity seen in both age groups (Fig. 1B), yet we did not formally explore this pattern further and instead focused on testing the differences between conditions as our target analysis.

Planned comparisons showed that, in adults, dwell time to the face was significantly higher in pretense than in real scenarios for one simple and the two complex actions (Action ‘*eating a cookie*’: Mean (SE), M_real_ = 37.4% (2.2), M_pretend_ = 48.7% (2.1), t(173) = 3.69, P < 0.001; Action ‘*serving and eating jelly*’: M_real_ = 23.8% (2.3), M_pretend_ = 38.8% (2.1), t(180) = 4.86, P < 0.001; Action ‘*serving and drinking juice*’: M_real_ = 13.5% (2.2), M_pretend_ = 23.4% (2.1), t(174) = 3.25, P = 0.001). For toddlers, while we observed higher dwell estimates in pretense, in particular for the action ‘*serving and eating jelly*’ (M_real_ = 23.8% (2.6) M_pretend_ = 30.7% (2.7), these differences were not significant (t(183) = 1.83, P = 0.069). All other comparisons were not reliable (P > 0.2). (to see raw mean values per group instead of estimated marginal means see *Figure S1* in SI).

Dwelling to the active hand (Fig. 1B) was analyzed using the same statistical approach as before, revealing a significant main effect of type of action (F(3,715) = 302.51, P < 0.001), and an interaction between type of action and condition (F(3,715) = 15.80, P < 0.001), with no other significant parameters. Pairwise comparisons between conditions showed similar results for both age groups. For the simple action *’eating a cookie*’, dwelling to the active hand was significantly higher in the real than the pretend movies at both ages (Toddlers: M_real_ = 17.6% (0.9), M_pretend_ = 15.0% (0.9), t(413) = −1.97, P = 0.050; Adults: M_real_ = 17.1% (0.8), M_pretend_ = 13.5% (0.7), t(389) = −3.44, P = 0.001), while for the complex actions, dwell time was reliably higher in pretend than in real at both ages for the action ‘*serving and eating jelly*’ (Toddlers: M_real_ = 0.8% (0.9), M_pretend_ = 4.1% (0.9), t(419) = 2.50, P = 0.013; Adults: M_real_ = 0.4% (0.8), M_pretend_ = 3.5% (0.7), t(408) = 2.93, P = 0.004) and only in adults for the action ‘*serving and drinking juice*’ (M_real_ = 6.5% (0.8), M_pretend_ = 9.2% (0.7), t(391) = 2.51, P = 0.012).

Along with total looking time, which provides an aggregated measure of attention, we measured the number of fixations within each AOI, defined as looking instances for at least 100 ms (ref. 27), more related to exploratory behaviors (Fig. 1A, middle row). Following the same modeling approach as above, in this case with the number of fixations to the face as the outcome variable, we found a significant main effect of age (F(1,105) = 5.86, P = 0.017), condition (F(1,105) = 6.19, P = 0.014) and type of action (F(3,714) = 52.69, P < 0.001), which were qualified by a significant two-way interaction between age and type of action (F(3,714) = 2.85, P = 0.036), condition and type of action (F(3,714) = 21.15, P < 0.001) and a marginally significant triple interaction between age, condition and type of action (F(3,714) = 2.55, P = 0.055).

Planned comparisons showed significantly more fixations to the face in pretense than in real only for the complex actions in both age groups (Action: ‘*serving and eating jelly*’, Toddlers: M_real_ = 3.8 (0.5), M_pretend_ = 5.3 (0.5), t(202) = 2.14, P = 0.034; Adults: M_real_ = 3.8 (0.4), M_pretend_ = 6.7 (0.4), t(199) = 4.87, P < 0.001; Action: ‘*serving and drinking juice*’, Toddlers: M_real_ = 3.2 (0.5), M_pretend_ = 4.7 (0.5), t(199) = 2.09, P = 0.038; Adults: M_real_ = 4.4 (0.4), M_pretend_ = 6.9 (0.4), t(192) = 4.20, P < 0.001).

For the active hand, on the other hand, we observed a significant main effect of condition (F(1,106) = 7.64, P = 0.007) and type of action (F(3,717) = 255.20, P < 0.001), but also a significant two-way interaction between these factors (F(3,717) = 14.91, P < 0.001). Pairwise comparisons across actions showed significantly more fixations in pretense than real again only in the complex actions (Action: ‘*serving and eating jelly*’, Toddlers: M_real_ = 0.2 (0.2), M_pretend_ = 1.1 (0.2), t(396) = 2.80, P = 0.005; Adults: M_real_ = 0.2 (0.2), M_pretend_ = 1.0 (0.2), t(386) = 3.26, P = 0.001;. Action: ‘*serving and drinking juice*’, Toddlers: M_real_ = 2.2 (0.2), M_pretend_ = 3.4 (0.2), t(385) = 3.68, P < 0.001; Adults: M_real_ = 2.4 (0.2), M_pretend_ = 3.2 (0.2), t(369) = 3.39, P = 0.001).

As a further measure of attention, we measured the number of times observers visited the face from another AOI (Fig 1A, bottom row). Mixed linear modeling results for the revisits to the face revealed main effects of condition (F(1,104) = 10.12, P = 0.002) and type of action (F(3,713) = 18.39, P < 0.001), yet an interaction between these factors (F(3,713) = 7.94, P < 0.001) and between condition and type of action (F(3,713) = 17.46, P < 0.001). Similar to the number of fixations, planned comparisons showed significantly more visits to the face in pretense than in real condition for the complex actions in both age groups (Action: ‘*serving and eating jelly*’, Toddlers: M_real_ = 2.9 (0.3), M_pretend_ = 3.8 (0.3), t(262) = 2.37, P = 0.019; Adults: M_real_ = 2.6 (0.2), M_pretend_ = 4.0 (0.2), t(257) = 4.85, P < 0.001;. Action: ‘*serving and drinking juice*’, Toddlers: M_real_ = 2.5 (0.3), M_pretend_ = 3.5 (0.3), t(257) = 2.78, P = 0.006; Adults: M_real_ = 3.3 (0.2), M_pretend_ = 4.6 (0.2), t(246) = 4.67, P < 0.00).

For the active hand, main effects of condition (F(1,105) = 10.14, P = 0.002) and type of action (F(3,715) = 263.69, P < 0.001) were observed, with however a significant interaction between these factors (F(3,715) = 15.35, P < 0.001) and a marginally significant triple-way interaction between age, condition and type of action (F(3,715) = 2.59, P = 0.052). Pairwise comparisons showed significantly more visits to the active hand in pretense than in real for complex actions at both ages (Action: ‘*serving and eating jelly*’, Toddlers: M_real_ = 0.2 (0.2), M_pretend_ = 1.2 (0.2), t(371) = 3.37, P = 0.001; Adults: M_real_ = 0.2 (0.2), M_pretend_ = 1.1 (0.2), t(362) = 3.54, P < 0.001; Action: ‘*serving and drinking juice*’, Toddlers: M_real_ = 2.6 (0.2), M_pretend_ = 3.7 (0.2), t(361) = 4.11, P < 0.001; Adults: M_real_ = 2.6 (0.2), M_pretend_ = 3.3 (0.2), t(346) = 2.85, P = 0.005).

Together, these results show that both toddlers and adults allocated more attention to the face of an actress performing make-believe actions compared to real-world actions, as measured through more fixations and revisits to the face. Adult observers also displayed more overall (aggregated) looking at the faces in pretend, and in general, the effects were seen for the complex actions. The active hand was also differentially attended in pretense, again mostly in pretense actions, and exhibiting similar profiles in toddlers and adults.

### Gaze shifts

Along with measuring where subjects attend, dynamic AOI analysis can also tell us how attention switches between visual regions via gaze shifts (rapid eye movements) between AOIs (see **Figure 2A**). Gaze shifts, according to computational accounts^32^, index how observers sample and integrate visual information over time. As in the previous section, we kept our focus on evaluating shifts between the face and the active hand and employed the same statistical approach as before, but here we fitted a generalized linear mixed model of shift counts (with the same fixed and random factors as above) using a Poisson distribution to model the number of shifts counts. Results are shown in **Figure 2B** (for a display of individual raw means estimates see *Figure S3* in SI).

**Figure 2.**
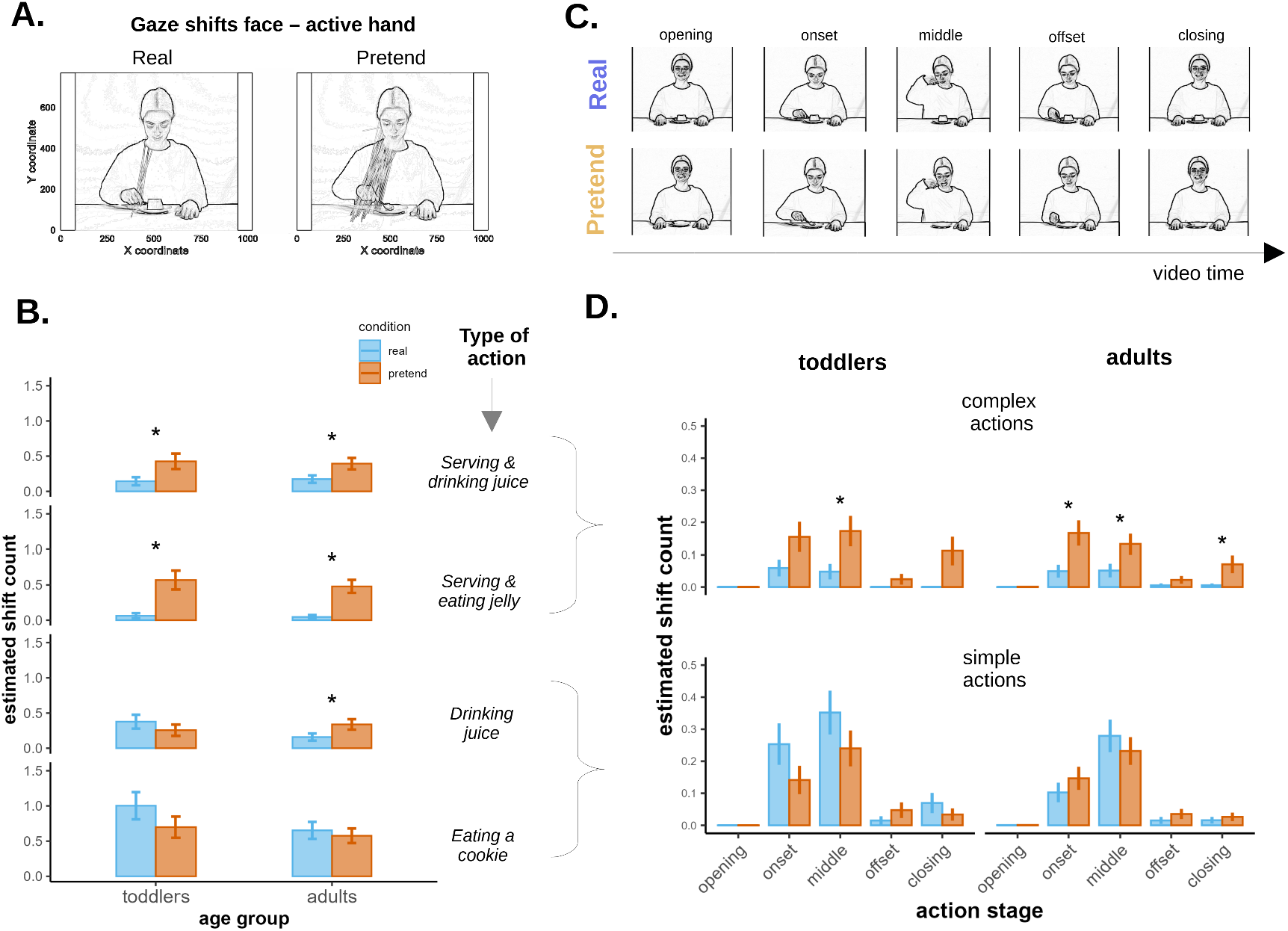
Gaze shifts between the face and the active hand. **A**, snapshots of the action ‘*serving and eating jelly*’ with all gaze shifts between the face and the active hand measured for this action across age groups overlaid as straight lines (for the same plots with all the actions see *Figure S2*). **B**, estimated marginal mean (EMM) of shift counts for each action type, arranged top-to-bottom from more complex to simpler ones. **C**, snapshots of the five stages the actions were divided in, depicted here for ‘*serving and eating jelly*’, but shared across actions *Opening*: the actress looks to the front smiling with both hands standing still over the table; *onset*: the actress moves her active hand towards the center of the table, grasps an object (spoon in this case) and serves a piece of jelly; *middle*: the actress moves her hand from the table towards the face, eats the jelly, and then moves the hand downwards; *offset*: she leaves the object over the table and returns the hand to its original position; *closing*: the actress looks again to the front and smiles. **D**, EMM of shift counts per action stage, combined for simple and complex actions, respectively. In B and D, error bars correspond to SE of the EEM counts *, P < 0.05.

We found a significant main effects of condition (χ^2^(1) = 18.36, P < 0.001) and type of action (χ^2^(3) = 90.11, P < 0.001), plus an interaction between these factors (χ^2^(3) = 39.10, P < 0.001). No further effects were significant (all Ps > 0.2). Planned comparisons showed that observers of both age groups performed significantly more shifts in pretense than in real actions for the complex actions (Action: ‘*serving and eating jelly*’, Toddlers: M_real_ = 0.06 (0.04), M_pretend_ = 0.57 (0.13), z = 3.50, P < 0.001; Adults: M_real_ = 0.05 (0.03), M_pretend_ = 0.48 (0.09), z = 3.8, P < 0.001; Action: ‘*serving and drinking juice*’, Toddlers: M_real_ = 0.14 (0.06), M_pretend_ = 0.43 (0.11), z = 2.30, P = 0.021; Adults: M_real_ = 0.17 (0.05), M_pretend_ = 0.39 (0.08), z = 2.22, P = 0.026).

Next, we asked *when* gaze dynamics differ between experimental groups, by decomposing the actions in successive stages (see **Figure 2C**). This allowed us to inquire, for instance, if gaze shifts between the face and the active hand accrue when the hand approaches the face of the actress (in the *middle* stage), thus ‘accompanying’ the hand movements, or emerge earlier in the action (in the *onset*, when the actress starts moving her hand), possibly reflecting an anticipatory behavior. **Figure 2D** displays these results. Here, we combined complex and simple actions to reduce the number of instances with zero shifts in each of the time stages evaluated. General inspection of results suggests that most shifts occurred during the *onset* and *middle* stages in both conditions and age groups. To statistically compare conditions, we employed the same modeling approach as for the global shift analysis, but fitting separate generalized mixed models for each stage.

Overall, we found that the differences between conditions were statistically reliable only for the complex actions. Specifically, for adults we observed more shifts during the stages *onset* (M_real_ = 0.05 (0.02), M_pretend_ = 0.17 (0.04), z = 2.65, P = 0.008), *middle* (M_real_ = 0.05 (0.02), M_pretend_ = 0.13 (0.03), z = 2.00, P = 0.045) and *closing* (M_real_ = 0.01 (0.01), M_pretend_ = 0.07 (0.03), z = 2.38, P = 0.017). For toddlers, we observed significant differences for the stage *middle* (M_real_ = 0.05 (0.02), M_pretend_ = 0.17 (0.05), z =2.25, P = 0.025). Note that although for the stage *closing* we observed more shifts in pretend than in real condition, statistical comparison between them could not be reliably done with the current model since the real condition did not contribute with shifts (it had all values with zero), causing the model’s estimated error ‘to crash’ for this comparison. However, testing the significance of the estimated shifts over zero for the pretend condition alone yielded significant results (estimate = 0.16 (0.05), t(20.24) = 3.52, P = 0.002), suggesting that the conditions do differ in the closing stage for toddlers too.

Thus, the analysis of gaze shifts over video stages provided a more nuanced characterization of gazing, showing that visual exploration departs between pretense and real actions right from the *onset* phase, i.e., when the actress starts moving her active hand horizontally to serve herself food or juice, onward. As for the attention measures reported in the previous section, these differences arose for complex actions. And together, they indicate that compared to literal actions, make-believe episodes induce more socially-oriented visual behaviors (social gazing) in mature and younger observers.

### Play mode and visual exploration

Along with testing the engagement of social processes, in our second goal we inquired whether pretense comprehension changes more broadly the manner in which subjects visually explore such scenes compared to real-world ones. We quantified how structured or predictable the exploration of visual areas was using gaze entropy (see **Figure 3A**), thus taking full advantage of the dynamic AOI approach, as it provided the trajectory of AOIs visited. Previous criteria for including trials in entropy analysis required instances having an amount of shifts among AOIs equal to or higher than the AOIs^35^. Here, simple actions had four AOIs (face, active hand, object and passive hand) and complex ones had five AOIs (the same as simple actions, plus another object). We included trials with a cut-off of 4 shifts across all actions (see Methods for further details).

**Figure 3.**
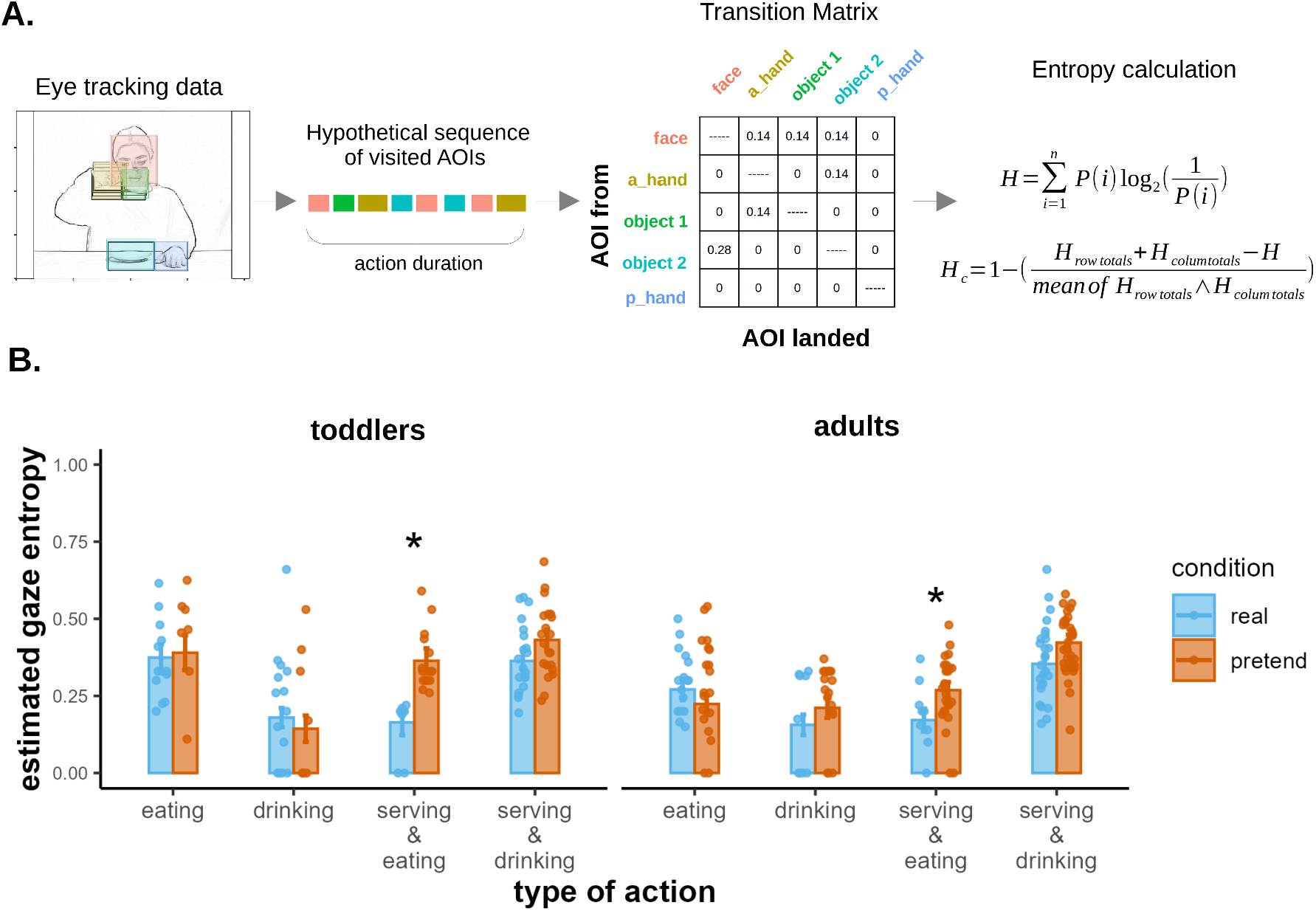
Structure during visual exploration of actions. **A**, main steps in the analysis of gaze entropy, a measure of order of the gaze signal^34^. For each stimulus presentation, gaze shifts among the AOIs are obtained from the eye tracking data, yielding a string of visited AOIs (illustrated in the middle of the panel); then, a transition matrix is derived with the probabilities (proportions) of shifts between each pair of AOI (excluding the diagonal, i.e., shifts within an AOIs). From this matrix, a measure of entropy is calculated using the formulas indicated on the right. The one below normalizes entropy values so that they range between 0 and 1 (the higher the values, the less structured the exploration) (see further details in Methods). **B**, estimated levels of entropy (marginal means) with SE as error bars; dots are the measured entropy values of included trials. *, P < 0.05.

To statistically compare the conditions (pretend, real) in each of the actions and age groups, we fitted entropy values to a generalized mixed effect model with the same fixed (age, condition, type of action) and random (intercept for subject) factors as the previous models, but now with beta distribution for the outcome variable, since entropy values are bounded between 0 and 1. Results are depicted in **Figure 3B**. Testing of model parameters yield significant main effects for the factor condition (χ^2^(1) = 5.94, P = 0.015) and type of action (χ^2^(3) = 83.30, P < 0.001), plus interactions between these factors (χ^2^(3) = 10.64, P = 0.014) and between age and type of action (χ^2^(3) = 9.09, P = 0.028).

Pairwise comparisons between conditions revealed significantly higher entropy levels in pretend in both age groups for the complex action ‘*serving and eating jelly*’ (Toddlers: M_real_ = 0.16 (0.04), M_pretend_ = 0.36 (0.04), z = 3.44, P = 0.001; Adults: M_real_ = 0.17 (0.04), M_pretend_ = 0.27 (0.03), z = 2.13, P = 0.033). Combining the simple and complex actions together, we found that for toddlers, complex pretend actions had higher entropy levels than real ones (z = 2.26, P = 0.024), whereas for adults the combined effect was not reliable (P > 0.1).

Taken together with the results of the social section, these findings indicate that not only the amount of visual information (as measured through gaze shifts and attentional patterns) but also the manner in which subject sample it is different in make-believe situations compared to real ones, supporting the idea that play-like scenarios change the way subjects explore the environment^31^, possibly through the induction of a decoupled cognitive state.

## Discussion

In this study, we tested the main hypothesis that pretense comprehension entails a social nature right from its onset in the second year and found consistent evidence supporting it, with observers of pretense gazing more frequently at the face of a person enacting pretense than observers of real-world actions. Furthermore, we could reveal a less structured pattern of visual exploration in make-believe scenarios, reinforcing the idea that a play mode changes the way visual information is gathered and processed. The striking similarity of outcomes between toddlers and adult observers can be understood as an effect of experimental replication, and supports developmental continuity of these processes.

The putative contribution of social processes in pretense development has been long inquired into^37^, and we seek to contribute to this debate by leveraging the capabilities of eye tracking dynamic techniques to provide direct evaluation of theories. Regarding adults’ results, consistent with our predictions and in line with brain studies^18–20^, we found that adult observers made more socially-oriented visual behaviors in nonliteral than in real scenarios, measured with three complementary measures of visual attention, namely, total looking time, number of fixations and number of revisits to the face of a person enacting pretend actions. The convergence of our findings with those from brain reports, showing activation of TOM regions in pretense comprehension, contributes to validating our approach of targeting the face as a proxy of underpinning social processes and reinforces the notion that adults engage such processes when understanding pretense actions.

In the case of toddlers, we found similar patterns of looking as in adults, again placing more attention to the face of the actress in pretense actions, measured by the number of fixations and revisits. Dwell time comparisons, however, were not reliable at this age. As this measure integrates global gaze properties over time, it may be less sensitive at detecting moment-to-moment changes in visual processing, compared to fixations and revisits. Studies indeed show that aggregated measures of visual exploration and fixations conform independent yet complementary measures of visual attention^38^. Note, however, as we remarked in Results, that the absence of a triple interaction between age, condition and type of action for the dwell measure suggests that the way condition and type of action interact was similar across age groups. On the other hand, in both age groups we found that the differential effects of looking at the face across measures consistently arose for the complex, not simple actions. Previous reports have shown that toddlers of ∼ 15- to 24-months perform better in comprehension tasks with simple actions than complex ones^,3^. In these studies, during a simple scenario subjects were presented with two dolls, one of which was transformed in a pretend way, e.g., by pouring imaginary powder; in complex ones, the two dolls were transformed but one of them was then returned to its original status, e.g., so both were made dirty with imaginary powder, but one was subsequently ‘cleaned’. In both instances, subjects had to describe the status of the transformed doll or select an ad-hoc action ‘to fix’ its situation. In the simple scenario, subjects needed to track the consequences of one action, while in complex ones, they needed to track chains of two actions. This is similar to our distinction if we consider the number of steps involved during the onset phase of the action. The fact that we observed more social gazing behaviors in complex scenarios may seem surprising given this evidence on performance differences. However, we believe that rather than a difference in the degree of comprehension afforded by toddlers in our study, the complex actions we presented, by having more steps, made it clearer or more salient to the observers the pretend nature of the acts, thus more effectively engaging the social gazing effects. Indeed, one of our complex actions (’*serving and drinking juice*’) was previously shown to be understood by toddlers of 15-months using looking time techniques^4,5^. Regardless of the specific cause for the effects of the type of actions, our results can be informative for future studies to help decide on which type of stimuli to use to more reliably elicit specific patterns of responses. Substantively, they revealed consistent visual explorations targeting the face in both age groups, providing direct empirical evidence indicating that pretense comprehension entails a form of social gazing from early on in development.

Along with measuring attention to the face, we inspected how observers looked to the active hand as action unfolded, given that attention to it has been described as important for action understanding^28^ and social interactions^27^. Across age groups, we found that observers attend more to the active hand in make-believe than in real scenarios, paralleling the results on the face, suggesting that the perception of it also plays a role in understanding pretense. To further prove this, we inspected the dynamic interchange of information perceived between the active hand and the face through gaze shifts, which again revealed more explorations in pretense actions. Interestingly, time course analysis helped us unpack that the bulk of the shift activity occurred during the onset and middle stages of the actions, indicating that gaze shifts may have tracked the movement of the hand as it approaches the face (in the middle stage), but also anticipated it, when the actress starts moving her hand in the horizontal plane of the table during the onset. Departures between conditions manifested earlier for adults, during the onset, and for both age groups were present in the middle and even at the end (closing stage) of the actions.

Previous studies have shown that the switching of attention between the face and the active hand of an interacting adult emerges in the first year of life and has been linked with joint attention^27^. In addition to this possibility, we think that the dynamic gazing to the face expresses some form of ‘query’ or sampling^32^ of social information, in line with proposals highlighting the face as a source from which to infer goals and intentions^25^, and consistent with studies showing that toddlers can use visual contact to ask for help under uncertainty^39^. From these observations, it is likely that in our context, the missing elements in the pretense scenes may have induced some uncertainty and information-seeking in the observers, triggering social gazing to infer the intention of the performer. This mindreading mechanism would then assist in grasping the non-serious, playful purpose of pretend actions, making them intelligible^10,11^. Critics of this metarepresentational interpretation^12^, however, have maintained that it is incomplete, for it does not explain how the possession and use of the concept *pretend* on the part of a child may support the understanding of the actions themselves. Such an overall understanding likely involves further recruitment of causal knowledge, as remarked recently^14^, which would help decode action goals and build expectations about outcomes in incomplete scenarios^2^ (a capacity already in place from at least the first year of life^39^). Thus, while the interplay between underpinning processes remains to be fully specified, the evidence gathered here and in the studies with older chidlren^16,17^ supports the engagement of social processes as an element of pretense. Interestingly, this socialization would not be tied to observers only, as 24-month-olds have been shown to gaze more often at an adult while themselves enact pretense compared to instrumental actions^41^, suggesting that pretend play may enhance bidirectional social exchanges and that toddlers interpret it as an inherently social activity.

But what level of awareness do observers have of their ongoing social inferences? One possibility is that metarepresentations (the representations of mental states of other people) are engaged in an implicit form, through an early emerging, non-verbal system that processes social information^42^. Alternatively, they may be instantiated more explicitly, consistent with brain reports^18–20^ indicating that in adults, pretense activates regions associated with explicit TOM tasks^21^. Inspection of our results, however, may suggest that although we observed parallels in gazing across age groups (supporting the idea of a shared underlying system between toddlers and adults), there were differences in the outcomes, with adults showing a more robust focus on the face as measured through aggregated looking time. Yet the time course of gazing showed quite similar profiles among adults and toddlers in the gaze between the face and the active hand, suggesting that the differences could be more of a degree than of a kind. In this case, the same system -albeit presumably with different levels of maturation and integration with other systems^43^-may sustain metarepresentations in both age groups. Future studies will help clarify the nature of the representations engaged in early pretense (and TOM alike^36^) and adjudicate between these possibilities.

The second goal we addressed in this work was to evaluate whether, in addition to changing the way subjects gather social information, pretense also changes the overall pattern of visual exploration of scenes. Our results support this idea, showing that the manner in which observers explore the scenes is different in pretense, being less structured than in real-world scenarios.

Different accounts have long related play with exploration (see review in ref.43), and in the case of pretense, there is evidence showing that framing abstract reasoning problems as make-believe situations improves performance in children^29,30^. These remarkable findings support the notion that pretense engages a cognitive state or mode that not only helps players keep reality and imagination separated but also transforms the way information is internally processed. Our results are in line with and significantly extend these observations, suggesting that this mode is operative early in life and it is capable of modulating the very process of perceptual sampling and its organization, extending all the way up to include body displacements^31^ and the construction of abstract situation models, as the referred reports above have revealed.

In turn, because of these connections with exploration, the nature of this ‘play state’ has been linked with curiosity^44^, while other accounts have emphasized that a distinctive aspect of it is the adoption of a positive *attitude toward the possible*^6^. According to Singer and Singer (1992) (ref.6), it is this attitude that allows players to detach from the literal world and ‘to see’ it and explore it from creative angles.

Interestingly, converging lines of evidence point to the Default-Mode-Network (DMN) as a possible biological substrate of this state and pretense more broadly. The DMN is a large-scale brain network composed of nodes from the parietal, temporal and frontal lobes, that in adults has been implicated in detached forms of cognition and abstraction, including autobiographical memory and mind-wandering, among others (see reviews in refs.45,46). Along with activation of TOM regions which belong to this network, studies of pretend play also report higher activity in the hippocampus in pretense compared to real actions comprehension^18,20^. The hippocampus is a node of the DMN, which beyond its well-documented role in memory, it has been involved in imagination^47^. More related to our findings, gaze entropy, as we measured it here, has been shown to be modulated by the integrity of the hippocampus^35^, and along with studies on gaze reinstatement^48.49^ and neuroanatomy^50^, they indicate that there are pathways connecting this region with the oculomotor system, and that the way the eyes scan the environment would be in part modulated by its operations. Developmental studies also indicate that the DMN and the hippocampus in particular, are functional from at least the second year in humans^51,52^. Thus, the DMN and its joint functionality supporting both social and visual exploratory behaviors, offers an encompassing framework in our context, but further studies are necessary to determine if our results can be explained by specific patterns of activity of the DMN.

It is worth noting that the effects on social and visual exploratory behaviors reported here were obtained by testing a particular form of pretense (imaginary or pantomime), thus additional research including other forms (e.g., object substitution, attribution of animacy, role play, see e.g., ref.17) is necessary before assessing their generalisability. Likewise, and as we alluded to above, to determine how the underpinnings here exposed (visual exploration and social gazing) interact with perceptual and memory processes that ‘fill in’ the missing elements of pretense scenes^3^ (for instance via pattern completion^48^ and causal reasoning^14^) and which together help observers construe the overall symbolic meaning of actions. These insights can translate to computational approaches of pretense^53^, and continue broadening our understanding of how many children grasp and enjoy pretense games^54^, but also why many have difficulties entering them^55^.

## Materials and Methods

### Participants

A group of 44 healthy toddlers (mean age = 19.56 months, age range = 16.26 - 20.00 months; females = 25) and 65 adults (mean age = 20.55 years, age range = 19 - 24 years, females = 53) were recruited to participate in the study. Toddlers were recruited from pediatric health centers in Santiago, Chile. All participants came from low- to middle-socioeconomic-status, monolingual Spanish-speaking families. Children were born full-term and exhibited typical developmental trajectories at the time of intervention. Exclusion criteria included diagnosed visual or auditory impairments, neurodevelopmental disorders, and chronic conditions associated with developmental delays. Children with histories of neonatal asphyxia, epilepsy, chromosomal abnormalities, or inborn errors of metabolism were excluded. Adults were college students with no motor or sensorial difficulties. Three additional toddlers to the sample were not included in the analyses due to fussiness. Our study was approved by the institutional review board of the [Institution name, masked for reviewing]. Parents provided written informed consent before their child participated.

### Materials

A series of 8 movies with an actress performing real-world and pretend actions were created. We included 4 actions, two categorized as simple: *eating a cookie*, *drinking juice*, and two as complex: *serving and eating jelly*, *serving and drinking juice*. Each of these actions was enacted in a serious way with actual food or juice or by making the pantomime of the actions with no food or liquid.

The structure of each action is divided into 5 stages (as depicted in Figure 2C), beginning with the actress looking to the front and smiling, having both her hands standing over the table (*opening* phase). In the *onset* phase, she looks down and moves her right hand toward the center of the table, picks up an object (e.g., a spoon, a jar) and uses it to grab food (jelly) or pour juice. In the *middle* stage, she raises her hand toward her mouth, eats or drinks, and returns her hand to the center. In the *offset* phase, she leaves the object in its original position and moves her hand from the center to the right side of the table. Finally, during the *closing*, she looks to the front and smiles as in the *opening*.

In the simple action ‘*eating a cookie*’, the actress did not use an object to pick up a cookie, only her hand, and for the simple action ‘*drinking juice’*, she directly grabs the cup during the onset phase. For the complex actions, during the onset, she first grabs a spoon or jar and then takes a piece of jelly or pours juice into a glass, respectively for *serving and eating jelly* and *serving and drinking juice*.

### Design

Each participant was randomly assigned to either real (toddlers, n = 22, adults, n = 31) or pretend (toddlers, n = 22, adults = 34) groups. The real group saw video clips of real scenes first and then the pretend ones, while the pretend group was exposed to the opposite order. Each action (eating jelly, drinking juice, etc), was presented 2 times, alternately, with the same order across participants. For completeness, after presenting the stimuli of either condition, we presented the stimuli of the other condition. As in theory of mind studies with a similar design^36^, we focused our analyses on comparing the conditions that participants saw first, to avoid so-called ‘carry over’ effects (one condition influences the other). Interleaved between the video stimuli, we presented images of objects whose analysis does not form part of this report. The average time between stimulus presentations was 3 s, with a range of 2-4 s. The total duration of the experiment was ∼8 min.

### Procedure

For toddler evaluations, upon arrival at the lab, participants and their caregivers were familiarized with the environment and the experimenters. During the evaluation, infants sat in their caregivers’ laps, in front of the monitor that displayed the movies. Caregivers’ eyes were covered with an opaque glass during the entire duration of the experiment to avoid biasing infants’ behavior. Upon finishing the experiment, caregivers completed a questionnaire with socio-demographic information and were reimbursed for the costs of transportation to the university. Adult participants were invited to sit in front of the monitor and were not given special instructions beyond watching the movies passively.

Eye movements were recorded throughout the experiment using an eye-tracker system (Tobii, 1750, 50 Hz), with a sampling rate of 50 Hz. The screen had a resolution of 1024 x 768 px.

### Data analysis

Pre-processing: eye tracking data files were exported to be used in the R environment for processing. First, we inspected for trackloss (gazes out of the screen) and blinks (missing intervals of < 200 ms), and discarded all instances of stimulus presentation with more than 50% of trackloss (similar to ref.38) or 40 % of blinks. Of the 872 trials presented (109 participants x 8 trials/participant), 37 did not pass the criteria (4.2%), with 24 corresponding to toddlers and 13 to adults. Two toddler participants did not pass the inclusion criteria in all trials due to inattention and were not counted in the referred sample of n = 44 and discarded from the analyses.

#### Measures

Dynamic area-of-interest (AOI) analysis: we performed this analysis as in previous studies of action understanding^28^. For each video, we defined 5 AOIs: active hand, passive hand, face, object 1 (placed in the center of the table: the plate with the food in ‘*eating a cookie*’ and in ‘*serving and eating jelly*’, the cup in ‘*drinking juice*’, and the glass in ‘*serving and drinking juice*’), object 2 (for complex actions: the spoon in ‘*serving and eating jelly*’ and the jar in ‘*serving and drinking juice*’).

We obtained the AOI’s center positions at each video frame of the videos using the software Blender^56^, which allows for extracting the position of objects in movies over time. We inspected the resulting trajectories to ensure that the tracked points corresponded with the center of the AOIs. To derive gaze measures, the time-series of the AOIs were aligned with the eye-tracking data. Gaze positions were marked as falling in either of the AOIs or background (i.e., gazes within the screen boundaries but that did not match an AOI). When two or more AOIs overlapped, as when the active hand approached the face region, we used the criteria of the ‘most recent’ one as the AOI selected (i.e., the active hand, as it ‘enters’ the region of the face), as in previous studies^28^.

With this information we calculated attentional measures: % *dwell time*, as the total time that the gaze landed in an AOI, divided by the total video time; number of fixations, as the sum of instances where looking time to an AOI was at least 100 ms in duration; number of revisits, as the sum of instances where an AOI was visited after being in another AOI. *Gaze shifts* were defined as movements of gaze between AOIs within 200 ms. Ref 28. used 100 ms as the cut-off criteria in their study with children and adults; here, we increase the interval, as infants may be slower in their movements (see *Figure S4* in SI for a distribution of shift counts as a function duration). Gaze shifts that resulted from one AOI entering the region of another were not counted as such; only actual movements of gazes were counted.

*Gaze entropy* measures were used to quantify the level of structure of visual exploration as in Refs.35,57, following the procedure described there and in Figure 3A. In the formulas presented in the figure, H is the total entropy of the transition matrix, and P(i) are the values within the cells (which contain the proportion of shifts between each of the AOIs). A correction and normalization is then applied to account for the total entropy directed at each AOI, expressed in entropy for rows and columns (see Refs. 34,56 for further details). The measure of normalized entropy H_c_ ranges from 0 to 1 (with higher values meaning higher gaze entropy, or less structured explorations). Since the beta distribution we used to statistically analyze entropy outcomes is bounded (0,1), we added or subtracted an offset of 0.01 to floor or ceiling values, respectively.

### Statistical Analysis

As mentioned in the Results section, we focused our analyses in comparing the real and pretend conditions for each action and age group, after fitting each outcome variable to the corresponding statistical models described. To fit the linear mixed models, we used the functions lmer() from the R package (‘lme4’), with the following equation:

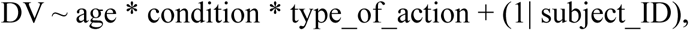

where DV stands for the dependent variables in Figure 1A at the trial level, age (toddlers, adults) and condition (real, pretend) were between-subject factors, type_of_action (with four levels, two simple and two complex actions) was a within-subject factor, and together with their interactions conformed the fixed factors in the models; the random component was captured in the intercept for subjects.

For the generalized mixed model used to analyze gaze shifts, we used the same model structure but with Poisson distribution, as it is appropriate for modeling count data, using the function glmer(). Entropy levels were modeled using a beta distribution for bounded data (0 to 1), using the function mixed_model() from the package ‘GLMMadaptive’. In all models, factors were summed coded to allow for testing of main effects, assessed with the functions anova() (for linear), Anova() (for generalized Poisson) and joint_test() (for generalized beta), all implementing type 3 sum of squares.

Extraction of estimated marginal means with their standard errors and pairwise comparisons were performed using the functions emmeans() and pairs() from the package ‘emmeans’.

## Author Contributions

C.G., C.J., C.M. and M.P. conceptualization and methodology; C.J., D.C. and M.P investigation; C.G. and C.M. and M.P. software/formal analysis; C.G. and M.P. writing.

## Conflict of Interest Statement

The authors declare no conflicts of interest.

## Acknowledgments

We would like to thank all the participants that contributed to the present work, including toddlers’ caregivers. We thank Consuelo De la Riva for their assistance in data collection and management. We also thank the National Center For Artificial Intelligence of Chile (CENIA), grant # FB210017, and the National Agency for Research and Development of Chile (ANID) for their financial support. C.J. acknowledges support from the grant FONDECYT # 11230621, C.M received support from the grant FONDECYT # 1251611, and M.P. received support from the grants FONDECYT #1241946, FONDEF IT24I0139. C.G. & M.P. acknowledge the Center for the Well-Being and Development of Adolescents and Children in the Digital Age (BAND, CIN250046), Chile.

## Notes

### Competing Interest Statement

The authors have declared no competing interest.

### Summary of Updates

Correction of methodological information; sections in the Discussion and References updated; Figures with higher resolution; Supplemental files updated.

## References

1. D. S. Weisberg, Pretend Play, WIREs Cogn Sci, 6, 249–261 (2015).

2. A. S. Walker-Andrews, P. L. Harris, Children’s comprehension of pretend causal sequences, Dev. Psy., 29, 915–921 (1993).

3. A. S. Walker-Andrews, R. Kahana-Kalman, The understanding of pretence across the second year of life, Br. J. Dev. Psychol. 17, 523–536 (1999).

4. K. H. Onishi, R. Baillargeon, A. M. Leslie, 15-month-old infants detect violations in pretend scenarios, Acta Psychologica, 124, 106–128 (2007).

5. J. Tee, C. Dissanayake. Can 15-month-old infants understand pretence? An investigation using the ‘violation-of-expectation’ paradigm. Acta Psychologica 138, 316–321 (2011).

6. D.G. Singer, J.L. Singer, The house of Make-Believe: Children’s Play and the Developing Imagination. Cambridge, MA: Harvard University Press; (1990).

7. G.A. Bateson, A theory of play and fantasy, in Steps to an Ecology of Mind, G.A. Bateson Ed. (New York, NY: Chandler, 1955), pp. 177–193.

8. P.L. Harris, R.D. Kavanaugh, Young children’s understanding of pretense. Monographs of the Society for Research in Child Development, 58, 1–91 (1993).

9. D. Skolnick, P. Bloom, What does Batman think about SpongeBob? Children’s understanding of the fantasy/fantasy distinction. Cognition, 101, 9–18 (2006b).

10. A. M. Leslie, Pretense and Representation: The origins of “Theory of Mind“ Psychol. Res., 94, 412–426, (1987).

11. O. Friedman, K.R. Neary, C.L. Burnstein, A.M. Leslie, Is young children’s recognition of pretense metarepresentational or merely behavioral? Evidence from 2- and 3-year-olds’ understanding of pretend sounds and speech, Cognition, 115, 314–319 (2010).

12. S. Stich S, J. Tarzia, The pretense debate. Cognition, 143,1–12 (2015).

13. J. Wolf, Implications of pretend play for Theory of Mind research. Synthese 200, 523, (2022).

14. D. Sobel, Understanding pretense as causal inference, Developmental Review, 68, 101065 (2023).

15. J. Perner, Understanding the Representational Mind, MIT Press (1991).

16. A.S. Lillard, R.D. Kavanaugh, The contribution of symbolic skills to the development of an explicit theory of mind. Child Dev., 85, 1535–1551 (2014).

17. M., Nielsen, & C. Dissanayake,. An investigation of pretend play, mental state terms and false belief understanding: In search of a metarepresentational link. Br. J. Dev. Psychol, 18, 609–624 (2000).

18. T. P. German, J. L. Niehaus, M. P. Roarty, B. Giesbrecht, M. B. Miller, Neural correlates of detecting pretense: Automatic engagement of the intentional stance under covert conditions, J. Cogn. Dev., 16, 1805–1817 (2004).

19. E. D. Smith, Z. A. Englander, A. S. Lillard, J. P. Morris, Cortical mechanisms of pretense observation, Soc. Neurosci., 8, 356–368 (2013).

20. C. Whitehead, J. L. Marchant, D. Craik, C. D. Frith, Neural correlates of observing pretend play in which one object is represented as another, SCAN, 4, 369–378 (2009).

21. C. Grosse-Wiesmann, A. D. Friederici, T. Singer, N. Steinbeis, Two systems for thinking about others’ thoughts in the developing brain, PNAS, 117, 6928–6935 (2020).

22. S. Kim, S. Kristen-Antonow, B. Sodian, A longitudinal study of early pretense: Metarepresentational or not. International Journal of Behavioral Development, 45, 345–354 (2021).

23. Y. Moriguchi, M. Ban, H. Osanai, I. Uchiyama, Relationship between implicit false belief understanding and role play: Longitudinal study. Eur. J. Dev. Psychol., 15, 172–183 (2018).

24. K. Libertus, A. Needham, Reaching experience increases face preference in 3-month-old infants. Dev. Sci.,14,1355–64 (2011).

25. A. Senju, G. Csibra, Gaze following in human infants depends on communicative signals, Current Biology, 18, 668–671 (2008).

26. Grossmann, T. (2010). The development of emotion perception in face and voice during infancy. Restor. Neurol. Neurosci., 28, 219–236 (2010).

27. S. Amano, E. Kezuka, A. Yamamoto, Infant shifting attention from an adult’s face to an adult’s hand: a precursor of joint attention, Infant Behav. Dev., 27, 64–80 (2004).

28. O. Ossmy, D. Han, B.E. Kaplan, M. Xu, C. Bianco, et al. Children do not distinguish efficient from inefficient actions during observation. Sci. Rep., 11, 18106 (2021).

29. M.G. Dias, P.L. Harris, The effect of make-believe play on deductive reasoning. Br. J. Dev. Psychol. 6, 207–221, (1988).

30. A. Wente, A. Gopnik, F.M. Fernández, T. Garcia, D. Buchsbaum, Causal learning, counterfactual reasoning and pretend play: a cross-cultural comparison of Peruvian, mixed- and low-socioeconomic status U.S. children, Phil. Trans. R. Soc. B, 377, 20210345 (2022).

31. J. Chu, L.E. Schulz, Not Playing by the rules: Exploratory play, rational action, and efficient search, Open Mind, 7, 294–317 (2023).

32. J. Gottlieb, Attention, learning, and the value of information, Neuron, 76, 281–295 (2012).

33. E. Risko, N.C. Anderson, S. Lanthier, A. Kingstone, Curious eyes: Individual differences in personality predict eye movement behavior in scene-viewing, Cognition, 122, 68–90 (2012).

34. N. Lee, V. Lazaro, J.J. Wang, H.H. Şen, K. Lucca, Exploring individual differences in infants’ looking preferences for impossible events: The Early Multidimensional Curiosity Scale. Front. Psychol. 13:1015649 (2023).

35. H.D. Lucas, M.C. Duff, N.J. Cohen, The Hippocampus promotes effective saccadic information gathering in humans, J. Cogn. Neurosci.. 31, 186–201 (2019).

36. T. Schuwerk, D. Kampis, N. Alessandroni et al. Action anticipation based on an agent’s epistemic state in toddlers and adults, PsyArXiv, 2025

37. L. Vigotsky, Play and Its Role in the Mental Development of the Child, Soviet psychology, 5(3), 6–18 (1968).

38. A. Zangrossi, G. Cona, M. Celli, M. Zorzi, M. Corbetta, Visual exploration dynamics are low-dimensional and driven by intrinsic factors. Commun. Biol., 4, 1100 (2021).

39. L. Goupil, M. Romand-Monnier, S. Kouider, Infants ask for help when they know they don’t know, Proc. Natl. Acad. Sci. U.S.A., 113, 3492–3496 (2016).

40. G. Csibra, S. Bíró, O. Koós, G. Gergely, One-year-old infants use teleological representations of actions productively, Cognitive Science, 27, 111–133 (2003).

41. H. Rakoczy, M. Tomasello, T. Striano, On tools and toys: How children learn to act on and pretend with ‘virgin objects’, Dev. Sci., 8, 57–73 (2005).

42. H. Rakoczy, The Development of Implicit Theory of Mind, in The Routledge Handbook of Philosophy and Implicit Cognition, J.R. Thompson, Ed. (Routledge, 2022), pp 336–350.

43. L. Wang, A.M. Leslie, Is implicit theory of mind the ‘real deal’? The own-belief/true-belief default in adults and young preschoolers. Mind & Language, 31, 147–176 (2016).

44. J. Chu, L. E. Schulz, Play, Curiosity, and Cognition Annu. Rev. Dev. Psychol., 2, 1–27 (2020).

45. J. Smallwood, B. C. Bernhardt, R. Leech, D. Bzdok, E. Jefferies, D. S. Margulies, The default mode network in cognition: a topographical perspective, Nat. Rev. Neurosci., 22, 503–513 (2021).

46. Y. Yeshurun, M. Nguyen, U. Hasson, The default mode network: where the idiosyncratic self meets the shared social world, Nat Rev. Neurosci., 22, 181–192 (2021).

47. A.E. Comrie, L.M. Frank, K. Kay, Imagination as a fundamental function of the hippocampus, Phil. Trans. R. Soc. B, 377, 20210336 (2022).

48. J.S. Wynn, J.D. Ryan, & B.R. Buchsbaum, Eye movements support behavioral pattern completion, Proc. Natl. Acad. Sci. U.S.A., 117, 6246–6254 (2020).

49. R. Johansson, M. Nyström, R. Dewhurst, M. Johansson, Eye-movement replay supports episodic remembering. Proc. R. Soc. B, 289, 20220964 (2022).

50. J.D. Ryan, K. Shen, Z.X Liu, The intersection between the oculomotor and hippocampal memory systems: empirical developments and clinical implications. Ann. N Y Acad. Sci., 1464, 115–141 (2020).

51. W. Gao, H. Zhub, K.S. Giovanello, J.K. Smith et al., Evidence on the emergence of the brain’ s default network from 2-week-old to 2-year-old healthy pediatric subjects, Proc. Natl. Acad. Sci. U.S.A., 106, 6790–6795 (2009).

52. T.S. Yates, J. Fel, D. Choi, et al. Hippocampal encoding of memories in human infants. Science, 387, 1316–1320 (2025).

53. L. Hengzhi, M. Tjandrasuwita, Y.R. Fung, A. Solar-Lezama, P. Liang, MimeQA: Towards Socially-Intelligent Nonverbal Foundation Models. 10.48550/arXiv.2502.16671 (2025).

54. A. S. Lillard, A. M. Pinkham, E. Smith, Pretend play and cognitive development, in Handbook of Childhood Cognitive Development. U. Goswami, ed., 2nd ed. (London: Blackwell, 2011), pp. 285–311.

55. F. J. Scott, S. Baron-cohen, ‘If pigs could fly’: A test of counterfactual reasoning and pretence in children with autism, Br. J. Dev. Psychol., 17, 349–362 (1999).

56. Blender Online Community (2022). Blender (Version 3.x) [Computer software]. Blender Foundation, Amsterdam. https://www.blender.org.

57. H.D. Lucas, A.M. Daugherty, E. McAuley, A.F. Kramer, N.J. Cohen, Dynamic interactions between memory and viewing behaviors: Insights from dyadic modeling of eye movements, J. Exp. Psychol. Hum. Percept. Perform., 49, 786–801 (2023).

