## Supplemental Information for "Pretend Comprehension Enhances Social and Exploratory Behaviors in Human Toddlers and Adults"

**This file includes:**

Supporting text
Figures S1 to S4
SI References

### Supporting Information Text

#### Rationale for reporting and displaying Estimated Marginal Means.

We deemed it appropriate to display estimated marginal means (EMM) instead of raw means, since EMMs account for the fact that some trials may have been missing, thus weighting accordingly the contributions of each subject and the associated error terms using all available data. Although in our case the % of missing trials was relatively low (4.2 %, as indicated in Materials and Methods), we still thought it was adequate to maintain this approach as in other reports[1]. For completeness, however, **Figure 1S** depicts the results of attention measures following the conventions used in Figure 1B but using raw means. Note that in this case, we did not perform statistical analyses for the comparisons between conditions with raw means (which were carried out using the outcomes of the mixed models described in the Main text and shown in Figure 1B).

#### Gaze shifts analyses: complementary information

**Figure 3S** plots the histogram of shift counts between the face and the active hand as a function of shift duration, for each condition and age group. Of the total of 700 shifts recorded, 599 fell below the cut-off criteria of 200 ms that we used (85.6 %). Nine of the total 700 shifts were very slow (> 1000 ms of duration) and are not displayed in the histogram.

### Figures

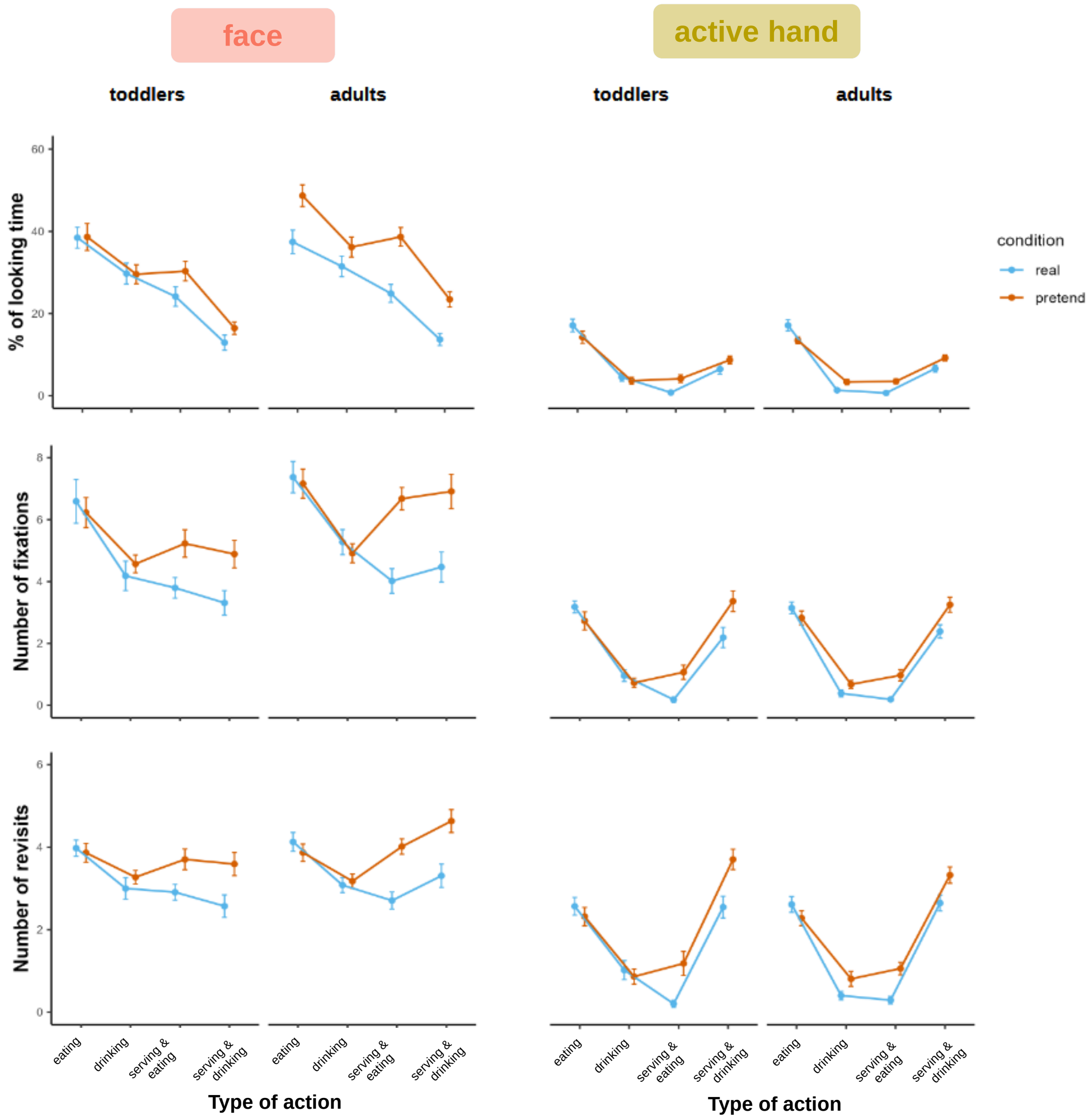


**Figure S1.** Raw means of attention measures derived from the dynamic AOI analysis. The organization is the same as in Figure 1B. Error bars correspond to standard errors.

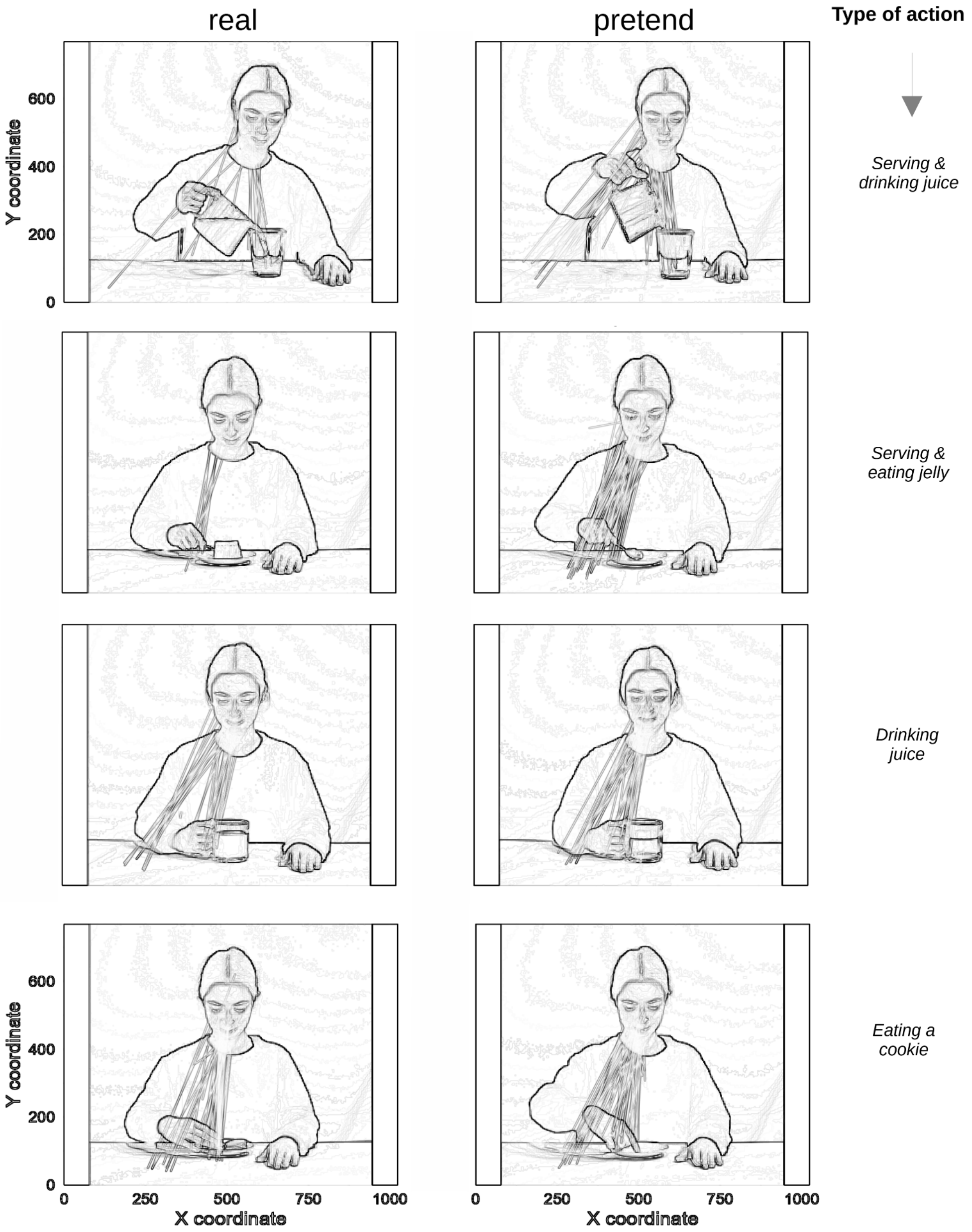


**Figure S2.** Snapshots of each movie action per condition with all gaze shifts between the face and the active hand across age groups, overlaid as straight lines (this figure complements Figure 2A in the main text).

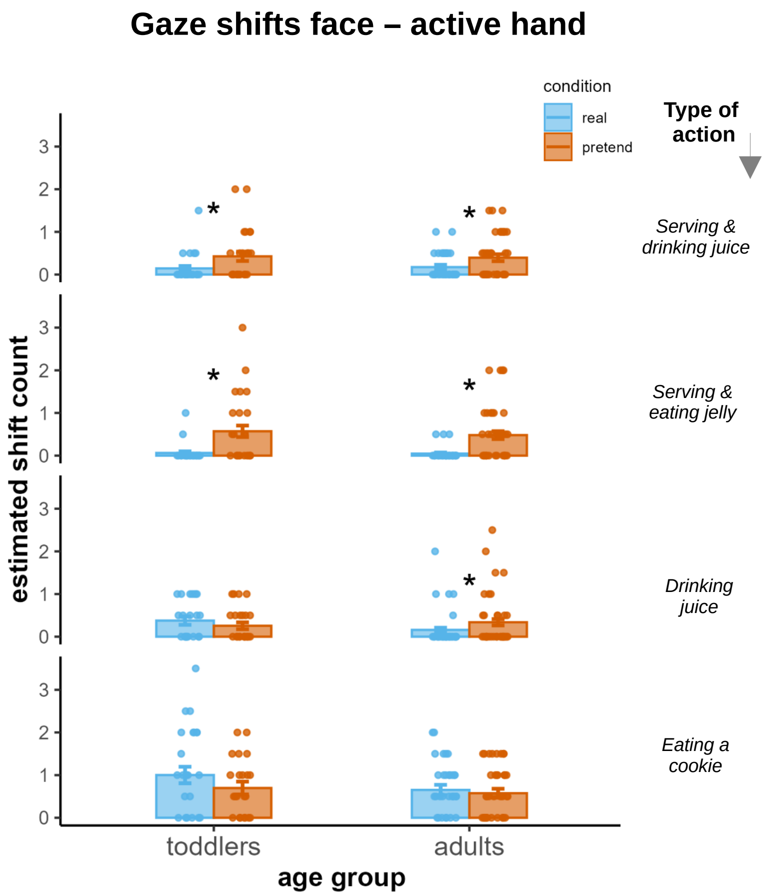


**Figure S3.** Estimated shift counts with subject's raw means overlaid as colored dots. Columns' height correspond to EMM values across subjects, resulting from the estimations of the generalized mixed model used (as described in the main text and Methods). *, P < 0.05 as in Figure 2B.

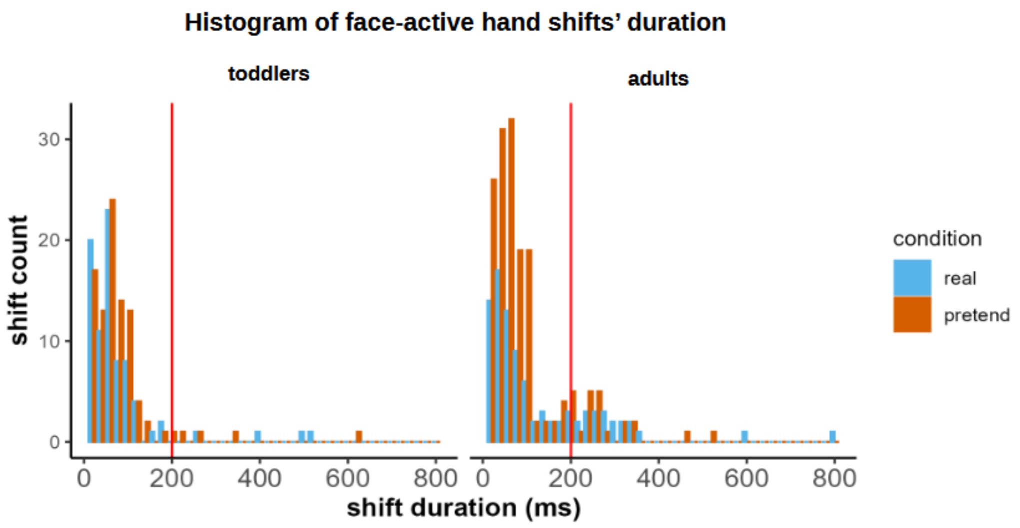


**Figure S4.** Histogram of shift counts over shift duration. The vertical red line corresponds to the cut-off value of 200 ms used in the analyses to be included as shifts.

### SI References

1. A. Wente, A. Gopnik, F.M. Fernández, T. Garcia, D. Buchsbaum, Causal learning, counterfactual reasoning and pretend play: a cross-cultural comparison of Peruvian, mixed- and low-socioeconomic status U.S. children, *Phil. Trans. R. Soc. B,* 377, 20210345 (2022).
